# Activation of M1 cholinergic receptors in mouse somatosensory cortex enhances information processing and improves detection behaviour

**DOI:** 10.1101/2023.06.06.543981

**Authors:** Wricha Mishra, Ehsan Kheradpezhouh, Ehsan Arabzadeh

**Author notes:** Correspondence: Ehsan Arabzadeh; Eccles Institute of Neuroscience, John Curtin School of Medical Research, The Australian National University, Acton, ACT 2601, Australia. These authors jointly supervised this work (Ehsan Kheradpezhouh, Ehsan Arabzadeh).

## Abstract

An important function of the brain is to form accurate representations of the world around us. To optimise sensory representations based on the demands of the environment, activity of cortical neurons is regulated by neuromodulators such as Acetylcholine (ACh). As such, ACh is implicated in cognitive functions including attention, arousal and sleep cycles. However, it is not clear how specific ACh receptors shape the baseline activity of cortical neurons and their evoked response to sensory stimuli. Here, we investigate the role of a densely expressed muscarinic ACh receptor 1 (M1) in information processing in the mouse primary somatosensory cortex (vS1) and in the animal’s sensitivity in detecting vibrotactile stimuli. We show that M1 activation significantly enhances the evoked response of vS1 neurons and the reversal of this enhancement by blocking M1. In addition, we demonstrate that M1 activation results in faster and more reliable neuronal responses, which is manifested by a significant reduction in response latencies and the trial-to-trial variability in neuronal activity. At the population level, M1 activation reduces the network synchrony and thus enhances the capacity of vS1 neurons in conveying sensory information. Consistent with the neuronal findings, we show that M1 activation significantly improves performances in a vibrotactile detection task. Overall, the M1-mediated enhancement in sensory efficiency reflects a multiplicative gain modulation at the neuronal level, resembling the changes observed during high attention states.

## Introduction

To survive, animals need to process the arriving sensory information differently depending on the context; while the sound of rustling leaves may not be important in the burrow, it could warn a mouse of an approaching predator in the open field. Neuromodulators such as Acetylcholine (ACh) provide one mechanism through which animals fine tune sensory processing to reflect the demands of the environment ^1,2^. Acting through various subtypes of ACh receptors (AChRs), the cholinergic system modulates information processing across different cortical areas influencing the animal’s behavioural state and attention level ^1,3,4^. Consistent with this function, activation of muscarinic AChRs (mAChRs) is known to increase neuronal excitability and enhance the response to relevant stimuli ^3–5^. Despite growing evidence on the effect of muscarinic neuromodulation in sensory processing, it is not clear how mAChRs affect the encoding of sensory inputs in cortical neurons and ultimately determine the perceptual responses to those inputs. Here, we combine pharmacological manipulations with *in vivo* electrophysiological, 2-Photon Calcium (Ca^2+^) imaging and behavioural studies, to characterise how activation of M1 receptors affects sensory information processing and behaviour.

We employed the mouse primary vibrissal somatosensory cortex (vS1) as it provides an optimal model to investigate neuronal coding due to its functional efficiency ^6^, structural organisation ^7^ and ecological relevance ^8,9^. We demonstrate that M1 activation enhances the sensory evoked responses in the mouse vS1 neurons through a multiplicative gain modulation. We also show an M1-induced reduction in the *first spike* latency and the trial-to-trial variability in the evoked responses. We further show that activating M1 induces desynchronisation in a subpopulation of neurons, reminiscent of attentive states ^10^. Finally, we show that consistent with our neuronal findings, M1 activation significantly improves the ability of the mice in detecting vibrotactile stimuli applied to the whiskers. Together, these results depict a key role for M1 receptors in sensory processing and behaviour.

## Results

### M1 activation enhances evoked responses in vS1 neurons

We first characterised the expression of M1 across layers of the vS1 cortex through immunostaining (Supplementary Fig. 1). Consistent with previous observations ^11^, we found that M1 is prominently expressed in layers 2/3 and 5. We further identified that M1 is highly co-localised in the excitatory neurons (Supplementary Fig. 1a; colocalisation with CaMKII; Pearson correlation co-efficient = 0.77; n=25). To determine the effect of M1 modulation on sensory processing, we first performed loose cell-attached recording (juxtacellular configuration, Pinault, 1996) under urethane anaesthesia. We recorded the activity of individual vS1 neurons during local activation or inhibition of M1 using a paired pipette method^13^ (Fig. 1a). We recorded and labelled vS1 neurons under continuous application of artificial cerebrospinal fluid (aCSF, control), M1 potentiator (agonist, Benzyl Quinolone Carboxylic acid, BQCA, 10 μM) or M1 specific inhibitor (Telenzepine Dihydrochloride, TD, 1 μM), while we stimulated the contralateral whiskers. The stimuli consisted of a brief vibration (20 ms in duration), which was presented at 5 different amplitudes (0-200 µm). We investigated the sensory evoked responses in the contralateral vS1 under these three conditions.

**Figure 1.**
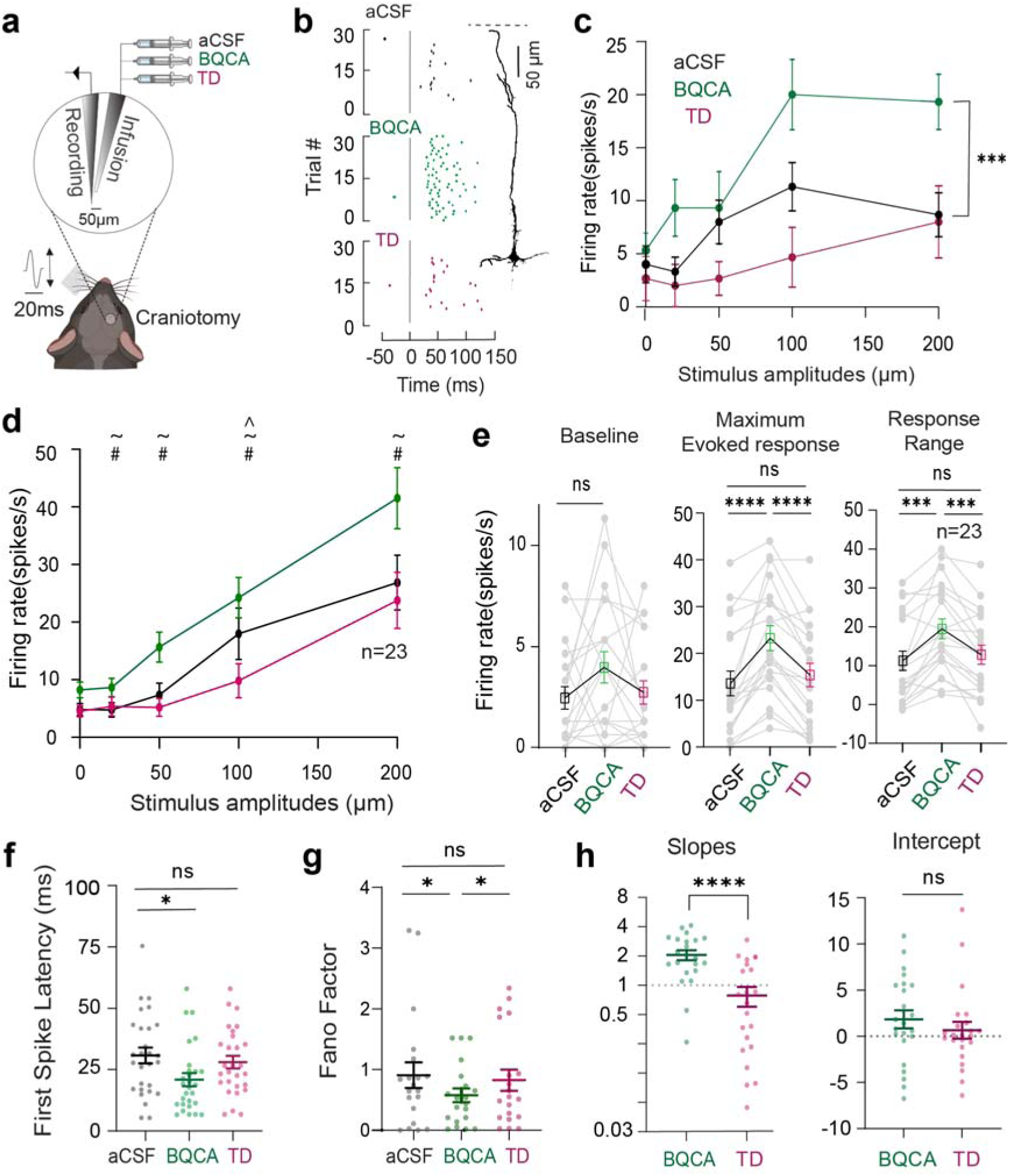
M1 activation enhances evoked responses in vS1 neurons. **a)** A schematic of the juxtacellular electrophysiology set-up for pharmacological manipulation of M1. The magnified circle depicts the custom-made pipette pair. **b)** Raster plots of spiking activity for an example neuron after application of aCSF (black, control), BQCA (green, M1 agonist) and TD (magenta, M1 antagonist). The grey vertical line represents the stimulus onset. The y-axis shows the trial numbers at subsequent presentations of the same stimulus. Inset: The reconstructed image shows the morphology of the example neuron, which is a layer 5 thick-tufted pyramidal neuron. The scale bar is 50 µm; the black dotted lines represent the dura. **c)** The input/output function of the neuron in **b)** across all conditions, with each dot representing the mean firing rate across 30 trials (p<0.0001, RM One-way ANOVA). **d**) The input/output function for all neurons under aCSF, BQCA and TD conditions. Each dot represents the mean firing rate across all neurons (n=23, RM One-way ANOVA, ∼-Significant statistical difference between BQCA and control; #-between BQCA and TD; ^- between TD and control. **e)** M1-mediated changes in baseline firing rate (left), evoked response (middle) and the response range (right). Grey dots indicate single neurons and the squares represent the mean values (n = 23). **f)** The first spike latencies calculated in a 100-ms window post stimulus presentation (200 µm). Each dot represents the mean latency of a neuron (n=23, p<0.05, RM One-way ANOVA). **g)** Fano factors of the evoked firing rate (200 µm, p<0.05, RM One-way ANOVA). **h)** Left: The slopes of the best fitted lines of neuronal response after M1 activation and inhibition. Each dot represents the slope of a neuron, and the black lines and bars represent the mean ± Standard error of mean (SEM) plotted on a logarithmic y-axis scale (n = 23, ****p<0.0001). Right: The y-intercepts of the best fitted lines (n = 23, p>0.05, Wilcoxon signed-rank test).

Figure 1b illustrates the effect of M1 modulation on the spiking activity in response to a 200-μm whisker deflection, recorded from an example neuron. M1 activation (BQCA) profoundly enhanced the stimulus-evoked responses, with no evident changes in the baseline activity (Fig. 1b, green). Subsequent application of M1 inhibitor (TD) reduced the evoked response of this neuron back to its initial level (Fig. 1b, magenta). Figure 1c illustrates how M1 modulated the response of the example neuron for the full range of stimulus amplitudes. Here, M1 activation produced an upward shift and M1 inhibition produced a downward shift in the response profile of the example neuron.

We observed a similar effect of M1 modulation across all recorded neurons (Fig. 1d; 200 µm, n = 23, p<0.001, RM one-way ANOVA). Figure 1e demonstrates the effect of M1 modulation on three main parameters of the neuronal response function: the baseline activity (amplitude = 0 μm), the maximum evoked response, and the response range (the difference between the maximum and minimum responses) as a measure of coding capacity. BQCA did not modulate the baseline activity (p=0.11, RM one-way ANOVA, Fig. 1e, left panel), but significantly increased the maximum evoked response (n=23, p<0.0001, RM one-way ANOVA, Fig. 1e, middle panel). Subsequent introduction of TD decreased the maximum evoked response to the initial values (n=23, p<0.0001, RM one-way ANOVA, Fig. 1e, middle panel). We observed a similar trend in the response range; BQCA significantly increased the response range (n=23, p<0.001, RM one-way ANOVA) and TD reduced it back to its initial values (Fig. 1e, right panel). These results indicate that M1 activation enhances the representation of vibrotactile inputs in vS1 neurons. In the following section, we investigate the effect of M1 modulation on other parameters of neuronal response including the latency and trial-to-trial variability.

### Temporal sharpening in vS1 neurons with M1 activation

The reliability of neuronal response and its timing can reflect the behavioural relevance of the stimulus; faster (reduced latency) and more reliable responses (less variability across presentations) suggest enhanced detectability at the neuronal level and ultimately better coding efficiency ^14,15^. On the other hand, higher variability in responses is detrimental to coding efficiency ^16^. The intrinsic variability in the response can be modulated by sensory stimuli ^17^ and non-sensory parameters including neuromodulation ^16,18^. Here, at the highest stimulus amplitude (200 µm deflection), we quantified the latency of the first evoked response. M1 activation with BQCA significantly reduced the first-spike latencies compared to the control condition Fig. 1f, n=23, p<0.05, RM one-way ANOVA); along with reduced jitter (Supplementary Fig. 2a, as seen by reduced Standard Deviation, p<0.001, Wilcoxon signed-rank test). The latencies increased by subsequent application of TD (Fig. 1f, p=0.178, RM one-way ANOVA).

To quantify the reliability of the evoked responses, we used Fano factor (the ratio of the variance to the mean of the firing rate) as a measure of trial-to-trial variability. A Fano Factor close to 1 reflects a Poisson distribution where the mean and variance of the response are the same ^19^. A higher Fano factor indicates a less reliable response from one trial to another (lower coding efficiency, Adibi et al., 2013). We applied this analysis to the evoked response at all stimulus amplitudes (Supplementary Fig. 2B). At the highest stimulus amplitude, M1 activation significantly decreased the mean Fano factor (Fig. 1g, p<0.05, Wilcoxon signed-rank test; aCSF versus BQCA); subsequent M1 inhibition increased the average factor (Fig. 1g, p<0.05, Wilcoxon signed-rank test; BQCA versus TD). These results show that M1 activation reduced the trial-to-trial variability in the evoked spike count among neurons, which indicates an increase in reliability. Using these data, we next investigate how these M1 modulations further affect the encoding of stimulus features.

### Multiplicative gain modulation in neuronal response function through M1 AChR

For every stimulus amplitude, we plotted the response under BQCA or TD condition against that response under aCSF condition; we calculated the slope and intercept of the best-fitted line for each neuron (Supplementary Fig. 2d). The slope and intercept provide information about the effect of M1 modulation on coding efficiency ^21^. The slope illustrates multiplicative changes in the neuronal response function and the y-intercept illustrates the additive changes. For example, a slope of 1 with a positive y-intercept would indicate an additive function signifying a consistent increase in response across all stimulus amplitudes. Conversely, a slope higher than 1 would indicate a multiplicative gain modulation signifying that the increase in activity is multiplicatively scaled from lowest to highest stimulus amplitudes.

Our results were consistent with an M1-induced multiplicative gain modulation in the response function. Overall, 91% of the recorded neurons exhibited a slope greater than 1 with a mean of 2.05±0.23 (Fig. 1h, green, p<0.0001, Wilcoxon signed-rank test). On the other hand, M1 inhibition reduced the gain of the response function, with an average slope lower than 1 (0.78±0.17, Fig. 1h, magenta, p<0.001, Wilcoxon signed-rank test). The y-intercepts did not show a systematic change with M1 activation or inhibition (p>0.05, Fig. 1h).

### Modulation of neuronal population activity by M1

We next determined the effect of M1 modulation on the population activity of cortical neurons through 2-photon Ca^2+^ imaging in both anaesthetised and awake mice. M1 is a G_q_-protein coupled receptor that increases intracellular Ca^2+^ through activation of various signalling cascades ^22^. Here, we expressed GCaMP7f (a highly sensitive Ca^2+^ sensor; ^23^) in layer 2/3 vS1 neurons. To modulate M1 activity, we implanted a cannula semi-parallel to the cranial window to perfuse the transfected area (see methods, Fig. 2a). The mice were allowed to recover for 4-5 weeks. Similar to previous experiments, we captured the effect of M1 modulation on the spontaneous activity and the evoked responses under aCSF (Control), BQCA (M1 activation) and TD (M1 inhibition) conditions. We then calculated changes in fluorescence (ΔF/F_0_) as a measure of neuronal activity.

**Figure 2:**
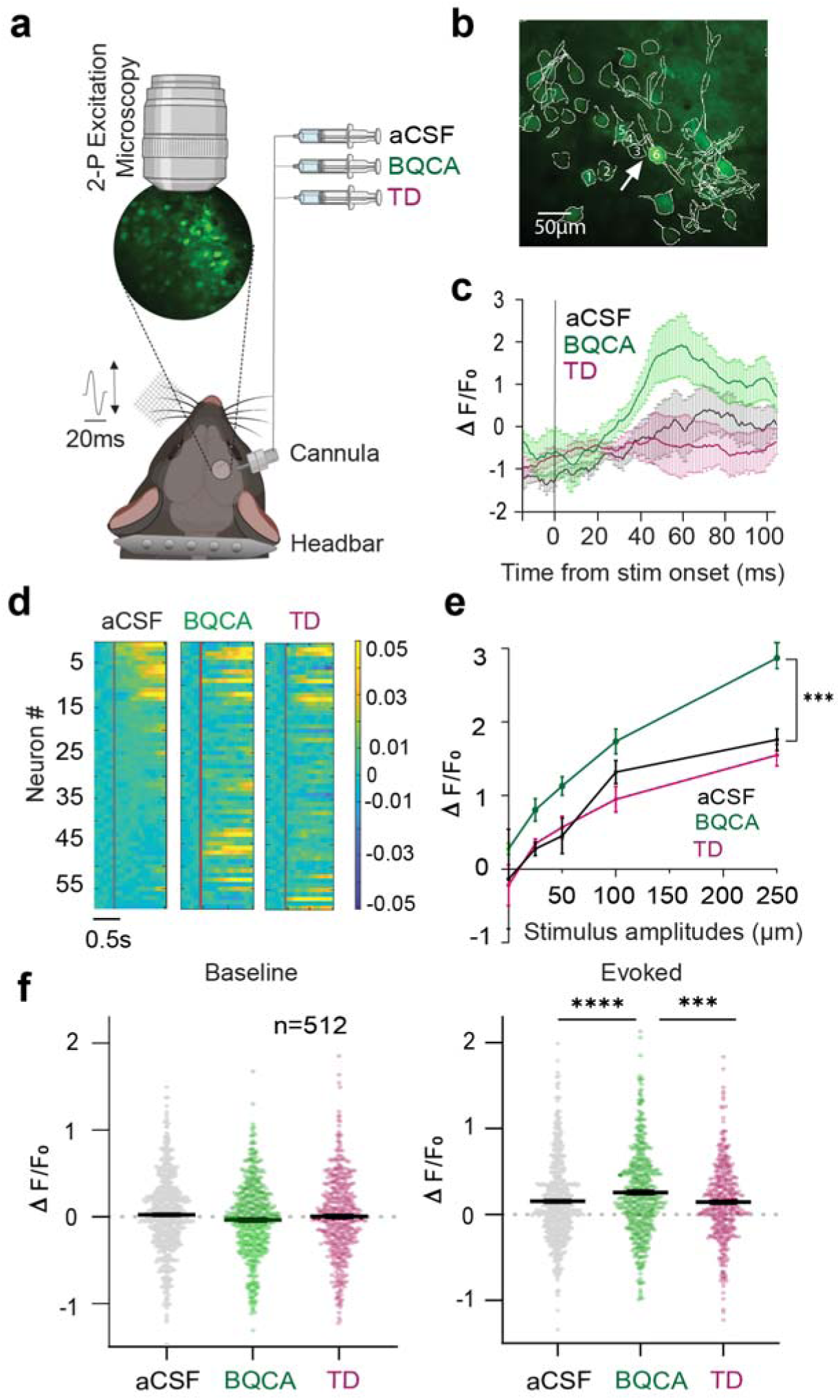
Characterisation of M1 modulation on neuronal population using 2-photon Calcium imaging. **a)** A schematic depicting the Ca^2+^ imaging setup for pharmacological manipulation of M1. The magnified circle depicts a 3mm cranial window expressing GCaMP7f in vS1. The changes in fluorescence activity were recorded from the neuronal population while applying vibrations to the contralateral whisker pad. **b)** A 2-photon image of layer 2/3 neurons expressed GCaMP7f after motion correction with ROIs highlighted. **c)** Changes in fluorescence (ΔF/F_0_) in response to the 250 µm whisker deflection, under the control (aCSF), M1 activation (BQCA) and M1 inhibition (TD) conditions for the neuron shown in **b** (white arrow). The grey vertical line indicates the stimulus onset. Shaded bars indicate standard error of the mean ΔF/F_0_ across 30 trials. **d)** A heatmap depicting the fluorescence activity of neurons shown in **b** (n = 60). The neurons in the aCSF (left panel) condition are sorted in a descending order and the sorting indices are conserved across BQCA (middle panel) and TD (right panel). Every horizontal line represents the ΔF/F_0_ of the same neuron across the three conditions, and the red line indicates the onset of a 250 µm whisker stimulation. **e)** Changes in ΔF/F_0_ measured in a 1 s window after stimulus with aCSF, M1 BQCA and TD across all stimulus amplitudes (n=512, 4 mice). The dots are the mean ΔF/F_0_; the error bars are the SEM. **f)** Baseline (left) and maximum evoked (right) neuronal response. Every dot represents a neuron. Black lines and error bars indicate the means and SEM. Here, data are pooled across four mice (n = 512, 4 mice). ***p < 0.001; ****p < 0.0001

Figure 2c captures ΔF/F_0_ in an example neuron (pointed with arrow, Fig. 2b) when the whiskers were stimulated at 250 µm amplitude in an awake head-fixed mouse. M1 activation enhanced the ΔF/F_0_ and subsequent M1 inhibition reduced the ΔF/F_0_ to its initial values. These findings were consistent across the neuronal population (same mouse, Fig. 2d, n=60).

We further quantified the effect of M1 modulation on neuronal population activity across all stimulus amplitudes. As observed in the example mouse, M1 activation significantly increased the sensory evoked response (Fig. 2e, n=512, 4 mice, p<0.01 for 50, 100 and 250 µm amplitudes); highest modulation was observed at 250 µm amplitude (Fig. 2f, right panel, n=512, 4 mice, p<0.001). Consistent with the electrophysiological data (Fig. 1e, right panel), we did not observe any significant modulation in the baseline activity (Fig. 2f, left panel, p=0.07, RM one-way ANOVA). Interestingly, we found a subpopulation of neurons, which remained silent (no evoked response) under the control condition (aCSF), but became significantly responsive to the stimuli after M1 activation (Fig. 2d, Neuron # 30-55). These results supported our previous findings that local activation of M1 in the vS1 cortex enhances neuronal excitability and sensory-evoked responses.

### Enhanced synchrony in vS1 neurons with M1 inhibition

We next investigated how M1 modulation affects the correlation of activity across recorded neurons. The capacity of a network to represent sensory information depends on the strength of the response to sensory stimuli (signals) as well as the similarity of responses in the absence of stimuli ^5^ (noise correlation). This similarity between stimulus-independent responses (noise correlations) can limit the encoding capacity ^14,24–26^ and cholinergic modulation has been shown to affect noise correlations ^5^.

Here, we investigated how stimulus-independent noise correlations are affected by M1 modulations. Figure 3a shows the fluorescence traces from 6 example neurons (Neurons 1-6, Fig. 3a) along with their correlograms (Neuron 5: Neuron 6, bottom) under the aCSF, BQCA and TD conditions. We observed high levels of correlation between example neuron pairs (most evident between Neuron 5 and 6) after M1 inhibition (Fig. 3a, bottom right) and this correlation was reduced by activating M1 (Fig. 3a, bottom centre). This finding generalised to all recorded pairs; M1 inhibition with TD showed a significant increase in pairwise correlation (Figs. 3b & 3c, p<0.01, 36 neuron pairs, Wilcoxon signed-rank test); and M1 activation with BQCA reduced pairwise correlations in the majority of the pairs (Fig. 3b, green, p<0.0001, 36 neuron pairs, Wilcoxon signed-rank test), resembling desynchronised activity. This trend could also be quantified across the neuron pairs shown in Figure 3A (Fig. 3c, p<0.0001, RM one-way ANOVA).

**Figure 3.**
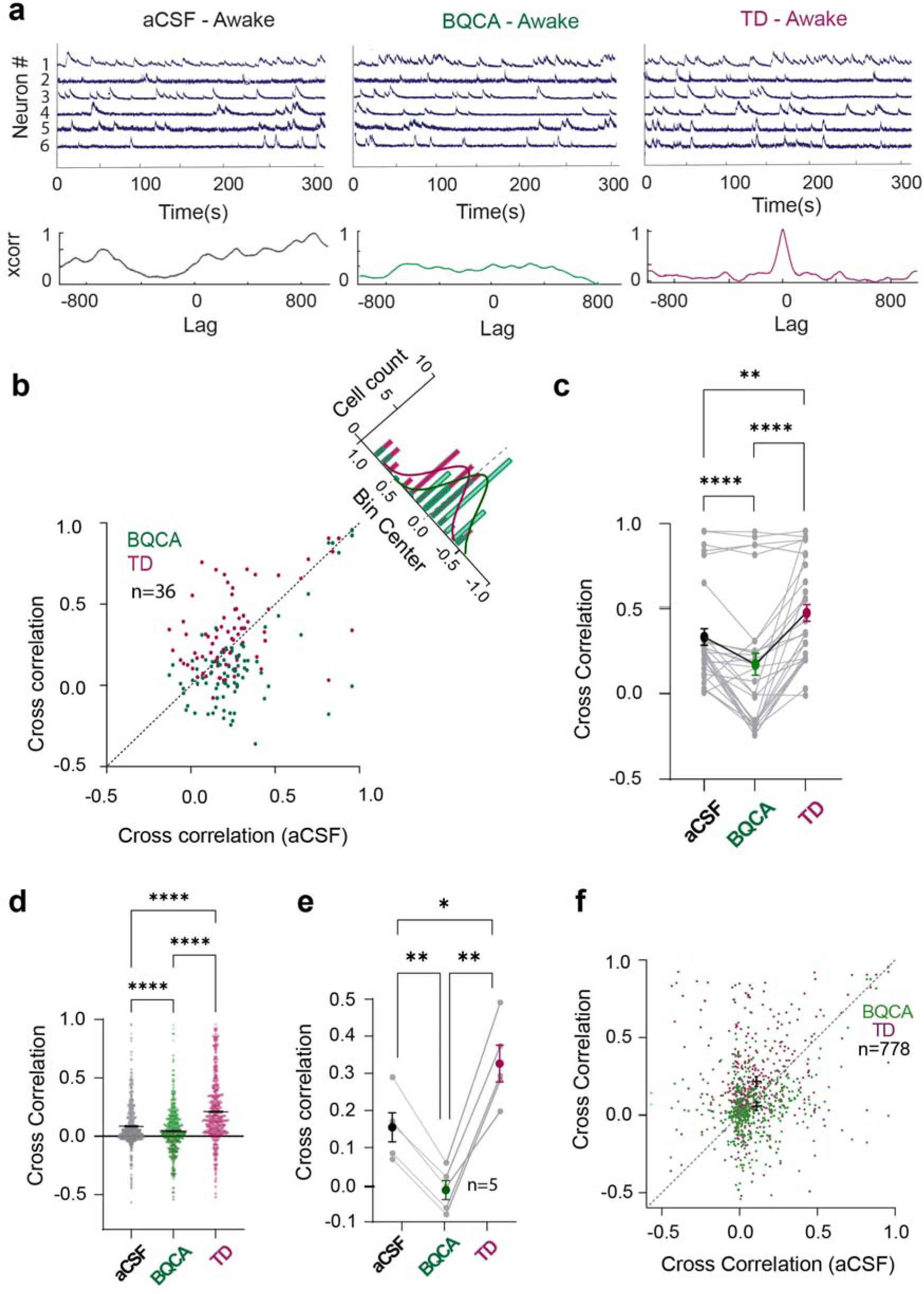
The effect of M1 modulation on neuronal synchrony. **a)** Raw fluorescence traces of 6 example neurons from one session in Fig. 2b over time under control (aCSF), M1 activation (BQCA) and M1 inhibition (TD) conditions. The spiking activity is more synchronised by inhibiting M1 (TD). The correlograms (below insets) show the cross correlation values versus lag for an example neuron pair (neuron 5: neuron 6). The sharp peak in the cross correlation values after TD perfusion indicates greatest synchrony between the example neuron pair at lag zero. **b)** Cross correlation of neuron pairs (for the 6 neurons shown in A, under the aCSF condition versus BQCA (green) or TD (magenta) condition. Neurons after M1 inhibition show an increased noise correlation as compared to M1 activation (p<0.0001, n=36, Wilcoxon signed-rank test). Inset: Histogram distribution for correlations after M1 activation (green) and M1 inhibition (magenta) from the line of equivalence. **c)** Average correlation coefficients in vS1 neurons in an example mouse with aCSF, BQCA and TD perfusions (n=36 neuron pairs, **p□<□0.01, ****p□<□0.0001, RM One-way ANOVA). Mean correlation coefficients are indicated by dots of black, green and magenta. **d)** Correlation coefficients across 5 mice with aCSF, BQCA and TD perfusions (n=778, 5 mice, ****p□<□0.0001, RM One-way ANOVA). Each dot represents the correlation coefficient of one neuron pair. The black bars indicate the mean correlation coefficients for each condition and the error bars represent SEM. **e)** Mean cross correlations in the aCSF, BQCA and TD conditions across mice (n=5, *p<0.05, **p<0.01, RM One-way ANOVA). **f)** Cross correlation across all animals and recording sessions, between aCSF, BQCA or TD. Neurons after M1 inhibition show increased noise correlation as compared to M1 activation (n=778, 5 mice). The dashed black line indicates the line of equivalence.

We applied this analysis to all responsive neurons among all animals and sessions; by activating M1, an enhanced desynchronisation was systematically observed (Fig. 3d, n=778, 5 mice, p<0.0001, RM one-way ANOVA). It is known that during desynchronised states, sensory information processing is enhanced both at the level of single neurons and in neuronal populations ^5,27–30^. Overall, these results support our previous findings that M1 activation improved information transmission across vS1 neurons. The enhancement in the sensory evoked response in vS1 neurons and the desynchronisation following M1 activation led us to investigate the effect of M1 modulation on mouse detection performance.

### M1 modulation enhances detection performance

To determine the effect of M1 modulation on detection behaviour, we tested a simple detection task in awake head fixed mice while modulating M1 activity. As nocturnal animals, mice regularly use their whiskers to navigate and explore their surroundings. Depending on the behavioural state of the animal (active engagement or quiet wakefulness), the efficiency of sensory information processing is altered by neuromodulatory inputs like ACh ^31^. Here, we investigated whether the observed enhancement in sensory processing through M1 receptors is reflected in the behavioural performance of mice.

We implanted a cannula in the right vS1 of 6 mice and attached a headbar on the back of the skull on Lambda to allow head fixation. After recovery, mice were trained to perform a whisker vibration detection task (Fig. 4a). Vibrations of different amplitudes were presented through a piezoelectric stimulator on the left whisker pad at amplitudes of 0, 15, 30, 60, or 120 μm. Mice received a sucrose reward for licking the spout on trials with vibration (15, 30, 60, and 120 μm) within a 400-ms window; licking in the absence of vibration (0 μm) was not rewarded (Fig. 4b). Stimuli were presented as blocks of 5 trials, containing 4 vibration amplitudes (15, 30, 60, and 120 μm) and a no-vibration trial (0 μm) in a pseudorandom order. This allowed us to calculate detection rates within each block (Methods, Behavioural analysis). To allow collection of a sufficient number of trials, only one solution was applied in a single behavioural session. As with earlier experiments, the solutions were aCSF (control), BQCA (10 µM, M1 activation) or TD (5 µM, M1 inhibition) and were perfused through the implanted cannula. These sessions were pseudo-randomly intermixed and each session was repeated 5 times. This produced an average 240 trials per condition (48 blocks X 5 trials). We found that the lick rate for 0-μm stimulus trials was similar to the pre-stimulus lick rate, indicating that mice successfully refrained from licking the spout in the absence of whisker vibrations (Fig. 4b; darkest line). As a general trend, mice licked at a higher rate and showed faster response times as the vibration amplitude increased (sample mouse, Fig. 4b, left inset, aCSF).

**Figure 4:**
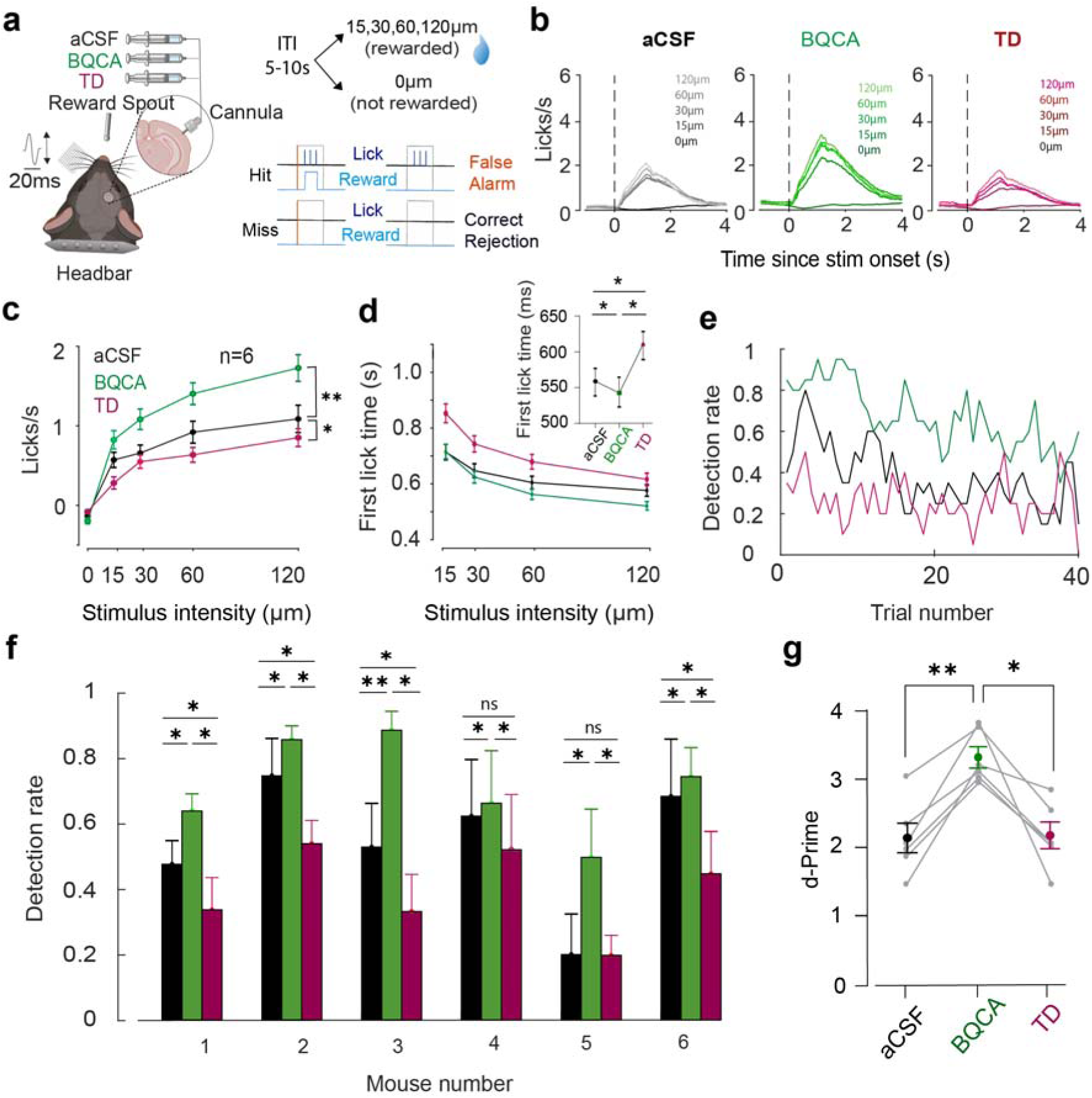
The effect of M1 modulation on mouse detection behaviour. **a)** A schematic depicting the behavioural paradigm. Inset: A 400-ms, 40-Hz vibration stimulus was presented at amplitudes: 0, 15, 30, 60, or 120 μm. Each stimulus presentation had an inter-trial interval of 5–10 s. **b**) The licking profile of an example animal showing the average lick rate against time, with the stimulus onset marked by the vertical black dotted line. Different stimulus amplitudes are depicted in different shades. **c)** The lick rates (n=6) across all stimulus amplitudes (0, 15, 30, 60, or 120 μm) in the control (black), M1 activation (BQCA, green) and M1 inhibition (TD, magenta) conditions. The solid dots represent the mean lick rates and the error bars represent SEM. **d)** Average first lick time (response time) across stimulus amplitudes for the control, M1 activation and M1 inhibition conditions. Inset: The first lick time across all 3 conditions for the highest stimulus amplitude of 120 µm (p<0.05, RM One-way ANOVA). **e)** Detection rate calculated across a session of 40 trials blocks (each block consists of 5 trials) for an example animal under the control, M1 activation (green) or M1 inhibition conditions (magenta). **f)** Average detection rate of the example animal in **E** under the control (black), M1 activation and M1 inhibition conditions. M1 activation significantly enhanced detection rate and inhibiting M1 decreased this detection back to baseline levels across animals. Mean detection rate is averaged over 5 sessions in each condition (p<0.05, RM One-way ANOVA). **g)** Mean d-prime across 6 mice: 2.088 (aCSF), 3.257 (BQCA), 2.125 (TD). Error bars meanLJ±LJSEM (**p<0.01, *p<0.05, RM-One way ANOVA).

Consistent with our findings at the neuronal level, we observed that BQCA improved the post stimulus lick rate (lick rates in sample mouse, Fig. 4b, middle inset, BQCA) whereas TD reduced the lick rate (sample mouse, Fig. 4b, right inset, TD). To better quantify the effect of M1 on detection, we compared the average response to the stimuli and the response time (the time of first lick after stimulus onset) among aCSF (control, black), BQCA (M1 activation, green) or TD (M1 inhibition, magenta) conditions (Fig. 4c). M1 activation significantly increased lick rates across all stimulus amplitudes (except 0), with the most significant rise observed at the highest amplitude (120 μm, green, Fig. 4c, n=6; p<0.01, RM one-way ANOVA). The lick rate decreased under TD condition even below the control rate (magenta, Fig. 4c, n=6; p<0.05, RM one-way ANOVA). In line with previous studies (Lee et al. 2020), we observed a reduction in the response time with increasing stimulus amplitude (Fig. 4d). The response (first lick) time for the highest stimulus amplitude decreased significantly after M1 activation (Fig. 4d, Inset, green; p<0.05, RM one-way ANOVA); M1 inhibition with TD increased this first lick time across all mice (Fig. 4d inset, magenta; p<0.05, RM one-way ANOVA).

To further investigate the effect of M1 modulation on perceptual sensitivity we quantified the detection rate for each mouse and condition (Methods, Behavioural analysis). As expected, the detection rates were generally higher at the beginning of a session (0-10 trials) and then gradually tapered towards the end of the session (Fig. 4e, black). As illustrated in the example mouse, activation of M1 produced a consistently high detection rate across all mice that was maintained for a higher number of trials (Figs. 4e & 4f, green) as compared to the control (Figs. 4e & 4f, black). The false alarm rates (response to no stimulus trials) were not significantly altered by M1 modulation (Supplementary Figs. 4a & 4b, p>0.05, RM One-way ANOVA), indicating that the enhanced detection rate is not a result of an overall increased lick rate. This increased sensitivity is directly captured in the d-prime measures (Fig. 4g). For every mouse, M1 activation (BQCA) enhanced the average performance (Fig. 4g; d-prime, aCSF: 2.08L±L0.21; d-prime, BQCA: 3.26L±L0.15) whereas, blocking M1 (TD) reduced the average performance to the control values (Fig. 4g, d-prime, TD: 2.125 ±L0.19). All together, these findings suggest that M1 activation improved perceptual sensitivity as was reflected in faster and more reliable responses.

## Discussion

To adjust the animal’s behavioural state to the demands of the environment, cortical activity is regulated by neuromodulators including the cholinergic system. Cholinergic input to the cortex has long been considered to act as a global activating system ^32,33^. In particular, layer 2/3 pyramidal neurons are powerfully influenced by Acetylcholine (ACh) through the dense projections they receive from the basal forebrain ^34^. Layer 2/3 is considered as a hub in cortical processing ^7,35^, where the majority of neurons fire sparsely due to a balanced feedforward excitation and feedback inhibition ^36^. ACh is thought to modify this balance to alter cortical activity and to shape the flow of information within the cortical circuits. Here, we showed that M1 activation significantly enhanced the sensory-evoked responses and reduced the trial-to-trial variability of these responses. This was illustrated by an increase in the signal strength and a decrease in the Fano Factors of the firing rates. At the population level, M1 activation reduced the network synchrony, which in turn enhanced the capacity of vS1 neurons in conveying sensory information. Consistent with the neuronal findings, we found that M1 activation improved performance in the vibrotactile detection task. Together, these findings show that M1 receptors enhance information processing in the somatosensory cortex and this is reflected in the animal’s ability to better detect sensory inputs.

Attention is known to dynamically change sensory representations in the cortex, with increased attention leading to an improved signal-to-noise ratio. The attentional modulations of sensory representations determine how we discriminate between stimuli ^37^ and integrate multiple sensory inputs ^38^. Previously, it has been shown that activating the cholinergic system enhances neuronal responses to sensory stimuli in a way that resemble a gain modulation ^39,40^. Such gain modulations reflect the changes in the sensitivity of a neuron to the stimulus while its selectivity for that particular stimulus is preserved ^41^ For example, neurons receiving a wide range of stimuli must be sensitive to weak stimuli but not saturated in response to stronger ones. In this way, neuronal responses can be continuously modulated to process sensory inputs at a wide dynamic range ^42^. Gain modulation causes an upward shift in the neuronal response function and strengthens representation of sensory stimuli ^4,43^. Here, we showed that M1 activation enhanced the evoked responses in vS1 cortex through a multiplicative gain modulation. Consistent with previous literature, activation of M1 led to changes in neuronal sensitivity, which in turn resulted in improved performance on sensory tasks by creating a more accurate representation of stimuli.

Cortical states are usually defined based on the correlated activity of neuronal populations ^27^. Several studies have reported enhanced sensory responses in desynchronized states due to lower noise correlations ^44,45^. During anaesthesia, sleep or quiet restful states, the cortex is in a deactivated state characterised by the presence of synchronised activity. In contrast, desynchronous firing is more prevalent during alert, attentive, and active behavioural conditions. Neuromodulators such as ACh can modulate changes in the dynamics of cortical activity, which can in turn shape the stimulus-dependent and stimulus-independent correlations ^5^. The stimulus-independent correlations between vS1 neurons, known as noise correlations, are typically higher in quiescent wakefulness compared to active exploration and whisking ^10,24^. In this study, we observed increased correlations between pairs of vS1 neurons when M1 was blocked; this is consistent with an increased synchronised activity with cholinergic inhibition ^5^. On the other hand, M1 activation induced desynchrony among neurons, indicating enhanced capacity for information coding at the population level. Previous studies have reported enhanced sensory evoked responses during desynchronised states due to reduced noise correlations ^44,45^. In line with this, our data showed that M1 activation enhanced whisker evoked responses (Figs. 1 & 2) and reduced synchrony at the population level (Fig. 3). This is consistent with studies highlighting that desynchronised states enhance response reliability in the somatosensory ^46^, visual ^5^ and auditory ^29^ cortices. Inhibitory interneurons play an important role in modulating this desynchronization ^47^. During quiet wakeful states, fast-spiking parvalbumin (PV) interneurons show synchronised firing ^48^ but are desynchronised during awake and attentive states. Somatostatin-expressing (SST) interneurons are also critical for precise synchronised firing in the vS1 ^49^. The excitability of PV and SST interneurons can be differentially regulated by muscarinic activation, with muscarinic activation exhibiting atypical hyperpolarising or biphasic responses in interneurons ^50,51^. Therefore, it is likely that M1 alters synchrony by differentially modulating the inhibitory drive in the vS1 neurons.

Cholinergic input to the cortex can vary dynamically depending on the level of arousal. For example, higher ACh levels are present in the cortex during awake attentive states as compared to lower ACh levels during quiet wakeful or anaesthetised states ^52,53^. The M1 potentiator, BQCA, used in this study increases the affinity of endogenous ACh to M1 receptors by binding to an allosteric site ^54^. Under anaesthesia, the M1-induced enhancement in the evoked response and baseline activity was less prominent as compared to the awake condition (Supplementary Fig. 3). These lower enhancements imply that M1-mediated modulations are potentiated to a greater extent during active awake states due to greater ACh released from the Basal Forebrain (BF). However, the presence of M1-mediated modulations during reduced cholinergic tone (i.e during anaesthetised states) suggests a fundamental role for M1 receptors in sensory processing. This is consistent with a recent study where cholinergic signalling through muscarinic activation facilitated auditory evoked activity in response to passive auditory stimuli, outside of any attentional context ^55^. Together, this indicates that a basal level of M1 activation plays an important role in passive sensory processing across sensory modalities. Therefore, it is likely that M1 modulates sensory processing through a unified mechanism that is preserved across sensory systems. It is interesting to note that the effects of M1 activation observed in this study - improved task performance, increased evoked response, improved neuronal response reliability and desynchronised firing – closely resemble the changes observed during enhanced attention ^43^.

The perceptual response to sensory input changes dynamically based on behavioural demands. Previous studies in rodents have shown that modulations in cortical state, such as those induced through muscarinic receptors, produce changes in behavioural performance ^26,56,57^. Based on this literature and as confirmed in our single cell recording and calcium imaging data, we predicted that modulations in M1 receptor activity would directly influence behavioural responses. We used the whisker vibration detection paradigm as an ideal model to study sensory processing due to the ecological relevance of the whisker pathway in rodent behaviour. Using their whiskers, rodents can be trained to learn complex behavioural tasks, such as discriminating textures ^9,58,59^, discriminating vibrations ^60,61^, and localising objects ^62,63^. In this study, we found that activating M1 produces an enhancement in the detection of vibrissal stimuli which was accompanied by a reduction in the response times (Fig. 4d). When an animal is actively engaged in the task, there is more cholinergic input into the vS1 from the basal forebrain ^31^ and applying BQCA to vS1 can potentiate the cholinergic response. In the visual system, muscarinic activation enhanced how the visual cortex responds to stimuli presented within an attended visual receptive field ^3^; and enhanced visual discrimination performance by engaging M1 ^64^. Muscarinic activation also facilitated auditory evoked responses in the auditory cortex ^55^. Our results are also consistent with a recent study that implemented an operant discrimination learning paradigm, where M1 inhibition reduced acquisition and consolidation ^65^. Together these findings suggest that M1 critically modulates behavioural performance across various modalities.

Despite the systematic findings on M1-induced neuronal gain modulation and enhanced behavioural responses, we observed some level of heterogeneity among neurons. Some responsive neurons showed reduced evoked response after M1 activation and some quiescent neurons exhibited an enhancement in evoked response after M1 inhibition (Fig. 2d, #Neuron 55-60). A cell-type specific modulation by M1 could explain this heterogeneity ^66^. M1 is predominantly expressed on cortical pyramidal neurons (Supplementary Fig. 1a) with a subset of inhibitory GABAergic PV interneurons (Supplementary Fig. 1b) and SST interneurons ^67^ also expressing M1. Parallels with the nicotinic system suggest that a small subset of neurons expressing a receptor can be very functionally relevant in feedback, feedforward or disinhibitory microcircuits ^66^. An important circuit motif for state modulation in the cortex is through disinhibition ^68,69^ consisting of PV, SST, and vasoactive intestinal peptide (VIP) interneurons. When VIP interneurons receive cholinergic projections from the BF ^70^, they remove the inhibition on layer 2/3 pyramidal neurons exerted by SST interneurons; thereby inducing a more active desynchronised state, similar to the M1-mediated desynchrony observed in this study. Therefore, it is likely that an M1-modulated disinhibitory microcircuit in layer 2/3 is responsible for this sensory sharpening (enhanced evoked response of excitatory pyramidal neurons). Future experiments will be necessary to determine how the cortical microcircuit is influenced by specific M1-mediated modulations of VIP, SST and PV interneurons during sensory processing.

## Acknowledgments

The experiments were supported by an Australian Research Council (ARC) Discovery Project (DP1701009), an NHMRC ideas grant (GNT1181643) and the ARC Centre of Excellence for Integrative Brain Function (ARC Centre Grant, CE140100007).

## Author contributions

W.M., E.K. and E.A. conceived and designed the project. W.M. performed the experiments. W.M., E.K., and E.A. analysed the data. W.M., E.K., and E.A. wrote the manuscript. All of the authors edited the manuscript and approved the final version.

## Resource availability

### Lead contact

Further information and requests for resources and reagents should be directed to and will be fulfilled by the Lead Contact, Wricha Mishra.

### Materials availability

This study did not generate new unique reagents.

### Data and code availability

The data generated in this study is available at: https://osf.io/rd8b4/. The MATLAB codes used in this study are available from lead contact on request.

## Methods

### SUBJECT AND EXPERIMENTAL METHODOLOGY

#### Mice

All experiments were performed on male and female C57Bl/6J mice (4-12 weeks old) housed in air-filtered and climate-controlled cages on a 12-12 hour dark/light reverse-cycle. All methods were performed in accordance with the protocol approved by the Animal Experimentation and Ethics Committee of the Australian National University (AEEC 2019/20 and 2022/16). Mice had access to food and water *ad libitum* except in behavioural experiments where mice were water restricted. The weight and overall health of all animals was monitored on a regular basis.

### METHOD DETAILS

#### Juxtacellular Electrophysiology

Mice were anaesthetised with a urethane/chlorprothixene anaesthesia (0.8 g/kg and 5 mg/kg, respectively) and placed on a heating blanket at 37°C. They were head-fixed on a custom-made apparatus. The scalp was opened via a 5 mm midline incision. After removing the scalp fascial tissue, a metal head plate was screwed to the posterior part of the skull and fixed in position with super glue and cemented subsequently. Once the cement had set, a 2 mm craniotomy was made above the right primary somatosensory cortex. The coordinates of the barrel cortex were marked as 1.8 mm posterior and 3.5 mm lateral to Bregma. The vasculature of the animal was also used as a reference to shortlist appropriate regions for recording.

Borosilicate glass pipettes were made by using a micropipette puller (P-97, Sutter Instruments) and custom-made programs. The recording pipettes had a tip diameter of ∼ 0.5-1 µm (impedance of 6-10 MΩ) and the infusion pipettes had a diameter of ∼20-30 μm with longer taper tips. The recording pipette was attached with glue to the infusion pipette on a custom-made stereotaxic setup with a tip-to-tip distance of 30-50 μm (Fig 1a, Kheradpezhouh et al., 2021).

The recording pipette was filled with a 2% neurobiotin (in Ringer’s solution). The infusion pipette was attached to a syringe pump (CMA402, Harvard Apparatus, Holliston, MA, USA) and filled with either artificial cerebrospinal fluid (aCSF), M1 receptor agonist Benzyl quinolone carboxylic acid (BQCA, 10 μM) or antagonist Telenzepine dihydrochloride (TD, 1 μM). The infusion pipette applied either aCSF, BQCA or TD at a flow rate of 2.5 μl/min. The pipette pair was positioned above the craniotomy and lowered using a micromanipulator. When the pipette pair reached the dura, 1 nA ON/OFF pulses (200 ms, 2.5 Hz) in current-clamp mode were applied. As the pipette touched the dura, Z-position of the micromanipulator was noted down for identifying the neuronal depth. The pressure in the recording pipette was maintained at 300 mmHg at this stage to avoid blockage of the pipette. After passing the dura, the pressure inside the recording pipette was reduced to 10-15 mm Hg, and the pipette was advanced at a speed of ∼2 μm/s while searching for neurons. The resistance was continuously monitored using the current clamp mode of a Dagan Amplifier (BVC-700A). Proximity to a neuron was observed by fluctuations in recording voltage and an increase in the resistance of the pipette (> 5-fold increase). At this step, the pressure was reduced to 0 mm Hg and juxtacellular (loose-cell attached) recording was performed. A custom-made MATLAB code provided the stimulus and recorded the neuronal response. Multiple recording session were made for all three conditions, aCSF, BQCA and TD. A total of 23 neurons were recorded from 19 mice.

At the end of the recording session, the recording pipette was moved closer to the neuron, which is indicated by an increase in the amplitude of voltage being measured (> 2mV). To further identify the morphology of a subset of neurons with neurobiotin by applying current pulses increasing in steps from 1 to 8 nA at a 200 ms duration. Successful loading was observed by broadening the AP spikes and a high frequency, tetanic like neuronal firing ^12,71,72^.

#### Vibrissal stimulation

A custom-made MATLAB code generated a pseudorandom sequence of stimulus amplitudes and acquired electrophysiological data through a data acquisition card (National Instruments, Austin, TX) at a sampling rate of 64 kHz. A wire mesh (2cm X 2.5cm) attached to a piezoelectric stimulator (Morgan Matroc, Bedford, OH) was slanted parallel to the animal’s left whisker pad (∼2mm from the surface of the snout) on the contralateral side, making sure that the whiskers reliably engage with the mesh. A consistent distance was maintained between the mesh and the face of the mouse. The whisker stimuli were composed of single Gaussian deflection amplitudes of 0, 25, 50, 100 and 200 µm for juxtacellular recordings, and 0, 25, 50, 100 and 250 µm for Calcium imaging. For behavioural experiments, the vibration stimulus was a train of discrete Gaussian deflections at amplitudes of 0, 15, 30, 60, or 120 μm. Each deflection lasted for 15 ms and was followed by a 10 ms pause before the next deflection.

#### GCaMP7f transfection and surgeries

Mice were briefly anesthetized with isoflurane (∼2% by volume in O2) and placed on a heating pad blanket (37°C, Physitemp Instruments). Isoflurane was passively applied through a nose mask at a flow rate of 0.4-0.6 L/min. The level of anaesthesia was monitored by the respiratory rate, and hind paw and corneal reflexes. The eyes were covered with a thin layer of Viscotears liquid gel (Alcon, UK). During this surgical procedure, the scalp of anaesthetised mice was opened along the midline using scissors and a 3-mm craniotomy was performed over the vS1 while keeping the dura intact. Expression of the calcium indicator GCaMP7f (Addgene, AAV1.Syn.GCaMP7f.WPRE.SV40) was achieved by stereotaxic injection of AAV virus. GCaMP7f was injected in the cortex at a depth of 230-250 µm from the dura at 4-6 sites (with four 32-nL injections per site separated by 2-5 minutes at the rate of 92 nLs^-1^). Following injections, a cranial window was covered using a 3 mm glass coverslip (0.1 mm thickness, Warner Instruments, CT). The animals were also implanted with a titanium headbar posterior to the cranial window, and a cannula (26 Gauge, Protech International Inc.) for microinjections of aCSF, BQCA or TD, immediately lateral to the cranial window. A small well was created around the cranial window using dental cement to allow water immersion for 2-Photon imaging. A thin layer of a silicon sealant (Kwik-Cast, World Precision Instruments, USA) was applied to cover all parts of the cranial window and skull.

#### 2-Photon Calcium imaging

3-4 weeks following the injection of GCaMP7f, the animal was transferred to a two-photon imaging microscope system (ThorLabs, MA) with a Cameleon (Coherent) TiLSapphire laser tuned at 920nm. The laser was focused onto layer 2/3 cortex through a 16x water-immersion objective lens (0.8NA, Nikon), and Ca^2+^ transients were obtained from neuronal populations at a resolution of 512 × 512 pixels (sampling rate, ∼30 Hz) (x16, 0.58NA). Laser power was adjusted between 40-75mW depending on GCaMP7f expression levels. All image acquisition was via ThorImage (ThorLabs, MA) and frames were synchronized with the stimulus presentation via the data acquisition card. In anaesthetised experiemnts, the animal was given an I.P. injection of urethane/chlorprothixene anaesthesia (0.8 g/kg and 5 mg/kg, respectively) and placed on a heating blanket. For awake recordings, mice were gradually habituated to the head-fixation apparatus – an acrylic tube with a custom-made headpost to allow head-fixation. After 3-4 days of habituation, mice were head-fixed in the apparatus and imaged. To study the effect of M1 modulation on neuronal response to whisker stimulation, aCSF, BQCA or TD was applied to this region of the cortex through the implanted cannula by switching between syringe pumps (CMA402, Harvard Apparatus, Holliston, MA, USA) at a speed of 2 µl/min. All videos were processed using the Python Suite2P package (https://github.com/cortex-lab/Suite2P) for motion correction and semi-automated ROI detection was performed in conjunction with ImageJ. The mean background neuropil was subtracted from each neuron’s calcium trace using a custom MATLAB script. The change in fluorescence (ΔF/F_0_) was quantified by using F_0_ as the mean fluorescence for each recording session.

#### Training and Behavioural Task

Mice implanted with the headbar and cannula were allowed to recover for 1 week, and placed on a water restriction schedule. The animals were gradually habituated to the experimenter and the head-fixation apparatus. The duration of placing the animal in the tube was increased gradually and once the animals were adequately habituated with the setup, they were held in position near the headpost with the help of homeostatic forceps, gradually increasing the duration of the hold. At each session, the mice were also presented with a 5% sucrose reward. Mice received unrestricted water for 2 hours immediately following the training sessions.

When the mice were well habituated to the setup, the first stage of training began where the animals received a reward for every lick. A vibration pulse (1s) followed each lick. This allowed the mice to lick reliably and get the sucrose reward. In the next stage of training, the mice were presented with a vibration till they licked the reward spout three times to claim the sucrose reward, after which a 60 s no-go period was enforced. In the last stage of training, the stimulus was either 120-µm (go) or 0-µm (no-go) with a variable inter-trial interval of 5-10 s. After mice learnt this version (above ∼85% correct), The vibration duration reduced from 1s in the first stage to 400 ms. The mice gradually learnt to lick the reward spout in response to a vibration of any amplitude (0, 20, 40, 80 or 120 µm). Stimulus amplitudes were pseudo randomised in blocks of 5 trials, with each block having all stimulus amplitudes.

A custom-made capacitive ‘lick-port’, connected to an Arduino UNO board (Duinotech Classic, Cat#XC4410), was used to deliver a sucrose reward and register licks. The lick-port was consistently positioned within reach of the mouth, ∼0.5 mm below the lower lip and ∼5 mm posterior to the animals’ snout. The capacitive voltage was sent to data acquisition card and a threshold determined the presence or absence of a lick.

#### Immunohistochemistry

At the end of the experiment, the animals were euthanised by an intraperitoneal injection of lethabarb (150 mg/kg). After opening the abdomen and chest medially, the heart was perfused with chilled normal saline followed by 4% paraformaldehyde in phosphate buffered saline (PBS) and the brain was harvested. The brain was fixed in 4% paraformaldehyde in PBS at 4°C overnight. After sequential rehydration with 10-30% Sucrose, the brain was sliced using a cryostat and incubated with streptavidin Alexa Fluor488 conjugate (Thermo Fisher Scientific, Waltham, MA, USA) overnight on a shaker at 4°C. For immunostaining of PV interneurons and pyramidal neurons (Supplementary Fig. 1), 100-µm thick coronal sections were permeabilised with PBS containing 1% Triton-X and 0.1% Tween 20 for 2-3 hours. To block non-specific binding sites slices were incubated in a blocking solution (0.25% Triton-X, 2% Bovine Serum Albumin in PBS), for 20-30 minutes at room temperature. Slices were then incubated with primary antibodies for M1 (Goat anti-M1 AChR, Abcam, Cat#ab77098, dilution 1:200), PV (Rabbit anti-PV, Abcam, Cat# ab11427, dilution 1:250) and CaMKII (Anti-CaMKII, Abcam, Cat#ab32678, dilution 1:250) added to blocking solution overnight at 4°C on a shaker. The following day, slices were washed and incubated with their respective secondary antibodies for 3-4 hours. Slices were then stained with 4′,6-diamidino-2-phenylindole (DAPI) to stain cell nuclei and mounted with Immu-Mount mountant (Thermo Scientific, Cat# 9990402) onto microscope slides.

### QUANTIFICATION AND STATISTICAL ANALYSIS

#### Neuronal Analysis

The spikes in each trial were extracted by applying a threshold for each neuron on the bandpass-filtered signal acquired during each recording session, using a custom-written MATLAB code. Neuronal firing rates were calculated by counting the number of spikes in each trial over a 50-ms window after the whisker-stimulus onset (0 ms). For every neuron and every stimulus amplitude and condition (aCSF, BQCA or TD), the mean firing rate (spikes/second) of 30 trials was reported. The latency of neuronal response was calculated as the timing of the first evoked spike in a 100-ms time bin, where the average firing rate was significantly higher than the baseline.

The Fano factor was calculated by dividing the variance (standard deviation squared) by the mean of the firing rate. Fano 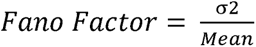

The best-fitting line, slope and intercept for each neuron were calculated and plotted using GraphPad Prism version 8.1.2.

#### Noise correlations

To calculate the noise correlation coefficient between neuron pairs, we computed the cross correlogram (using the MATLAB ‘xcorr’ function, and ‘coeff’ normalisation) of neuron pairs during periods of spontaneous activity in the absence of stimulus presentation. This allowed us to capture any stimulus independent correlations or noise correlations in neuronal activity. Cross correlation measurements were normalised to vary between 0-1. For each cell pair, the mean fluorescence activity (ΔF/F_0_) was correlated. The maximum height of the correlogram at lag 0 was taken as a measure of correlation strength.

#### Behavioural Analysis

Hit trials were defined as the presence of at least one lick 0-400 ms post stimulus onset and no licks 400 ms before stimulus onset and were used to calculate detection rate. The lick rate was calculated by subtracting the licks in a 500 ms pre-stimulus window from the licks in the post-stimulus reward window of 1s. To account for changes in motivation and engagement throughout the task, we excluded blocks of trials where the mice licked at 0-μm stimulus (false alarm). Here, the stimulus present trials (20, 40, 80 or 120 µm) were used to calculate the detection rate in each block (0 - No stimulus detected, 1 - All 4 stimulus intensities detected correctly). d-prime was computed for all trials by norminv (Hit rate) – norminv (False alarm rate), where norminv is the inverse of the cumulative normal function ^73^. Hits and False alarm rates were truncated between 0.01 and 0.99.

Relevant statistical analyses, p-values, and n-numbers are reported in figure legends and results section. Data were analysed and presented as mean ± standard error of the mean (SEM). Statistical significance was determined using MATLAB and GraphPad Prism version 8.1.2. A repeated measures (RM) one-way ANOVA with Bonferroni correction was used for analysing data which required a comparison of means across different conditions (aCSF, BQCA and TD) or a Wilcoxon rank-sum test for pairwise comparisons between 2 groups.

## Supplementary Figures

**Figure 1:**
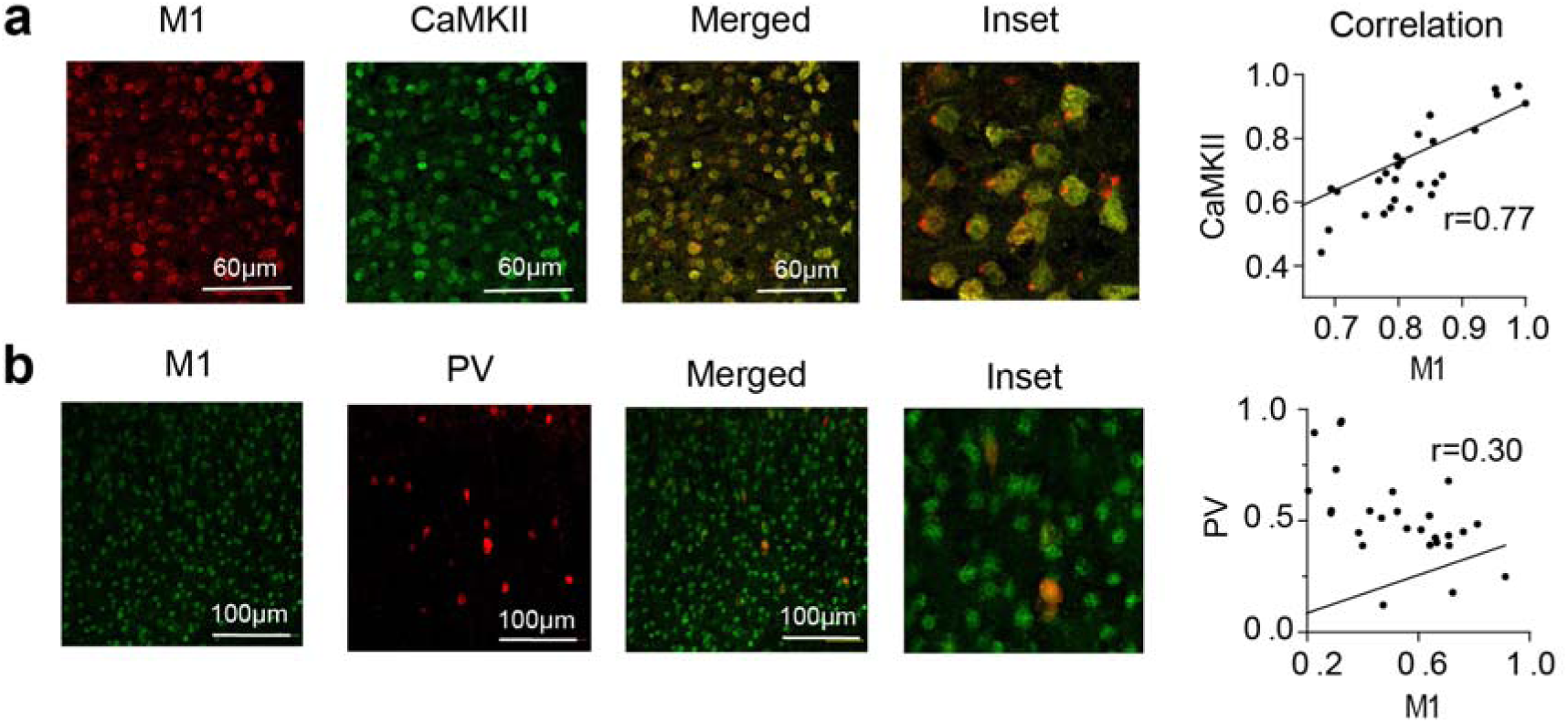
Expression of M1 on excitatory and inhibitory neuronal population in layer 2/3 of vS1. **a)** Co-localisation of M1 (Red) with CaMKII (green), a marker for excitatory neurons. M1 and CaMKII co-localise in a majority of neurons (inset). The scale bars are 60 µm. Pearson’s correlation coefficients for each neuron expressing CaMKII in a z-stack against the M1 expression (n=25). **b)** Same as a), but for M1 (green) and PV (red), a marker for an inhibitory interneuron subtype. Scale bars are 100 µm. Pearson’s correlation coefficients for each neuron expressing PV in a z-stack against the M1 expression (n=25).

**Figure 2:**
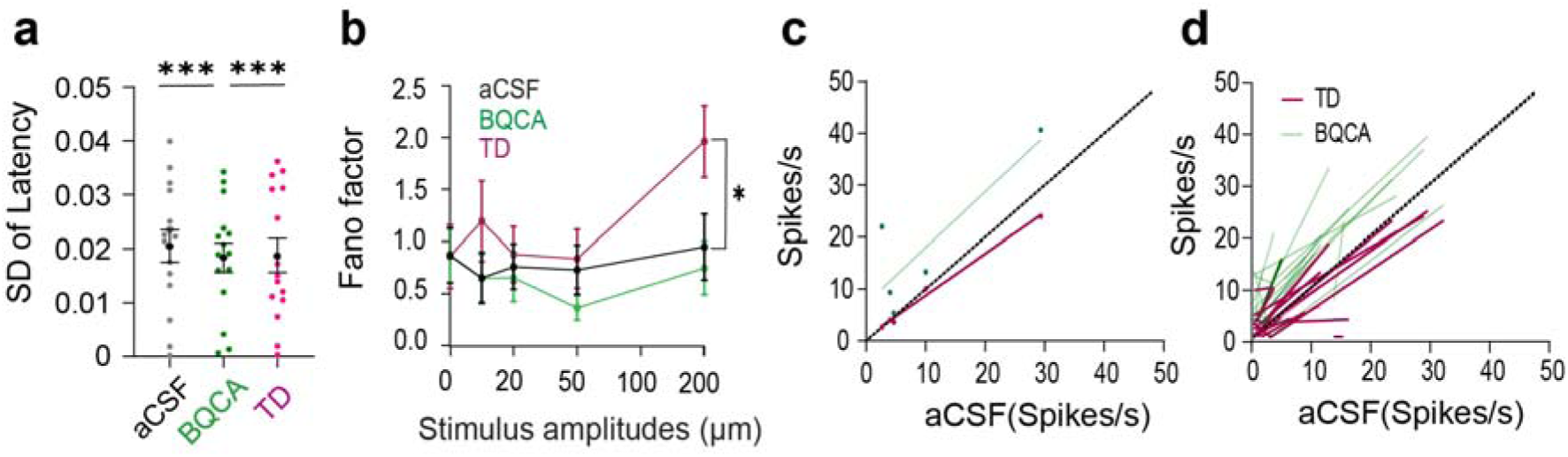
M1 activation enhances evoked responses in vS1 neurons. **A)** Standard deviation of first-spike latencies. n=23, ***p<0.001, RM One-way ANOVA. **B)** Fano factors of the spike counts for all stimulus amplitudes shown for aCSF (black), BQCA (green) and TD (magenta). The Fano factors for the largest stimulus amplitude (200 µm) increase significantly after applying TD, indicating increased variability. BQCA caused this Fano factor to decrease (*p<0.05, RM One-way ANOVA). **C)** The neuronal response of one example neuron after M1 activation (BQCA, green) and inhibition (TD, magenta) is plotted against the response in the aCSF condition. The green and the magenta lines illustrate the line of best fit used to calculate the slopes of the neuronal response; the black dotted line is the line of equivalence. **D)** The best fitted lines plotted for all neurons after M1 activation (BQCA, green) and inhibition (TD, magenta); the black dotted line is the line of equivalence.

**Figure 3:**
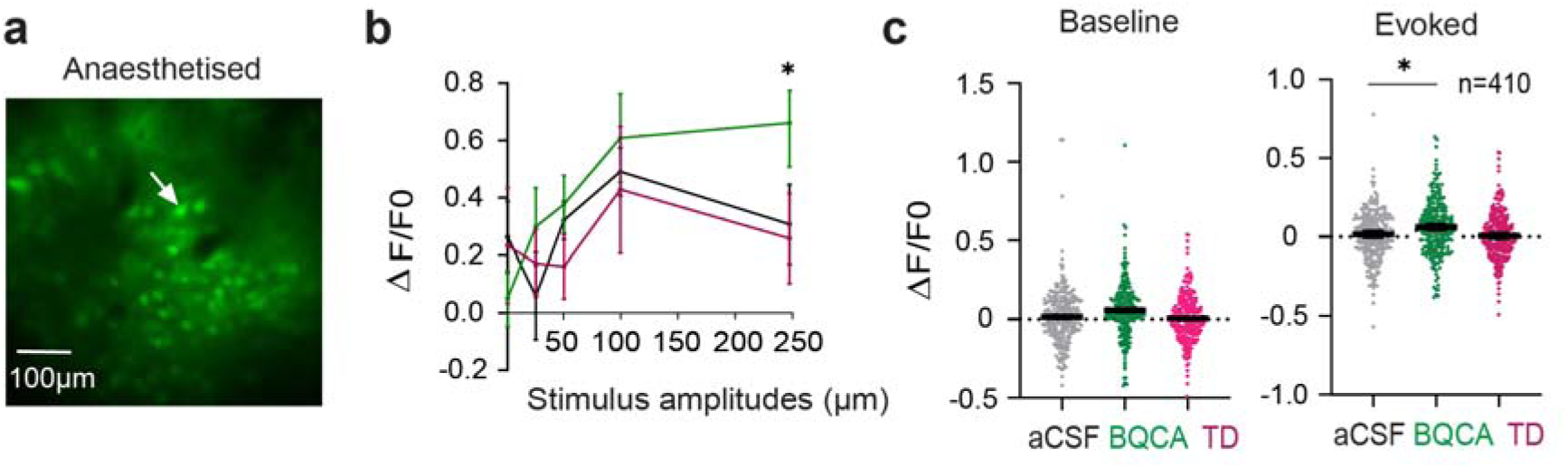
Effect of M1 activation on evoked responses in anaesthetised mice. **a)** An example plane of neurons imaged from layer 2/3 under urethane anaesthesia. **b)** Changes in fluorescence (ΔF/F_0_) in response to whisker vibrations, across the control (black), M1 activation (green) and M1 inhibition (magenta) conditions for the example neuron in A (white arrow). Error bars indicate standard error of the mean across 30 trials. **c)** Baseline and evoked neuronal response presented as ΔF/F_0_. Every dot represents a neuron. Black lines and error bars indicate the mean and SEM across neurons. Here data are pooled across four mice (n = 410, 4 mice). *p<0.05, RM One-way ANOVA.

**Figure 4:**
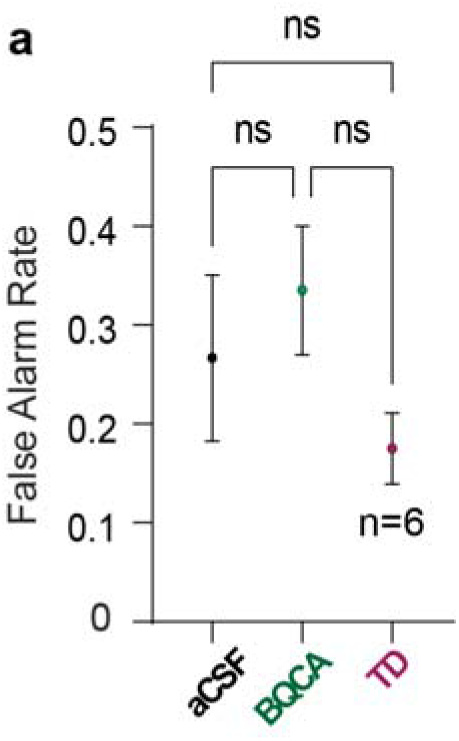
Effect of M1 modulation of False alarm rates. **a)** False Alarms across averaged across 6 mice, in the control (aCSF), M1 activation (BQCA) and M1 inhibition (TD) condition, shows no differences in False Alarm with M1 modulation. p > 0.05, RM One-way ANOVA.

## References

1. Parikh, V. & Sarter, M. Cholinergic mediation of attention: Contributions of phasic and tonic increases in prefrontal cholinergic activity. in Annals of the New York Academy of Sciences vol. 1129 225–235 (Blackwell Publishing Inc., 2008).

2. Sarter, M., Hasselmo, M. E., Bruno, J. P. & Givens, B. Unraveling the attentional functions of cortical cholinergic inputs: interactions between signal-driven and cognitive modulation of signal detection. Brain Res. Brain Res. Rev. 48, 98–111 (2005).

3. Herrero, J. L. et al. Acetylcholine contributes through muscarinic receptors to attentional modulation in V1. Nature 454, 1110–1114 (2008).

4. Herrero, J. L., Gieselmann, M. A. & Thiele, A. Muscarinic and nicotinic contribution to contrast sensitivity of macaque area V1 neurons. Front. Neural Circuits 11, (2017).

5. Minces, V., Pinto, L., Dan, Y. & Chiba, A. A. Cholinergic shaping of neural correlations. Proc. Natl. Acad. Sci. U. S. A. 114, 5725–5730 (2017).

6. Diamond, M. E. & Arabzadeh, E. Whisker sensory system – From receptor to decision. Prog. Neurobiol. 103, 28–40 (2013).

7. Feldmeyer, D. et al. Barrel cortex function. Prog. Neurobiol. 103, 3–27 (2013).

8. Brecht, M. & Sakmann, B. Dynamic representation of whisker deflection by synaptic potentials in spiny stellate and pyramidal cells in the barrels and septa of layer 4 rat somatosensory cortex. J. Physiol. 543, 49–70 (2002).

9. Petersen, C. C. H. The functional organization of the barrel cortex. Neuron vol. 56 339–355 (2007).

10. Harris, K. D. & Thiele, A. Cortical state and attention. Nat. Rev. Neurosci. 12, 509– 523 (2011).

11. Poorthuis, R. B., Enke, L. & Letzkus, J. J. Cholinergic circuit modulation through differential recruitment of neocortical interneuron types during behaviour. J. Physiol. 592, 4155–4164 (2014).

12. Pinault, D. A novel single-cell staining procedure performed in vivo under electrophysiological control: Morpho-functional features of juxtacellularly labeled thalamic cells and other central neurons with biocytin or Neurobiotin. J. Neurosci. Methods 65, 113–136 (1996).

13. Kheradpezhouh, E., Mishra, W. & Arabzadeh, E. A protocol for simultaneous in vivo juxtacellular electrophysiology and local pharmacological manipulation in mouse cortex. STAR Protoc. 2, 100317 (2021).

14. Averbeck, B. B., Latham, P. E. & Pouget, A. Neural correlations, population coding and computation. Nat. Rev. Neurosci. 2006 75 7, 358–366 (2006).

15. Peters, A. J., Chen, S. X. & Komiyama, T. Emergence of reproducible spatiotemporal activity during motor learning. Nat. 2014 5107504 510, 263–267 (2014).

16. Festa, D., Aschner, A., Davila, A., Kohn, A. & Coen-Cagli, R. Neuronal variability reflects probabilistic inference tuned to natural image statistics. Nat. Commun. 2021 121 12, 1–11 (2021).

17. Churchland, M. M. et al. Stimulus onset quenches neural variability: a widespread cortical phenomenon. Nat. Neurosci. 2010 133 13, 369–378 (2010).

18. Verhoef, B. E. & Maunsell, J. H. R. Attention-related changes in correlated neuronal activity arise fromnormalization mechanisms. Nat. Neurosci. 20, 969 (2017).

19. Charles, A. S., Horwitz, G. D. & Pillow, J. W. Dethroning the Fano Factor: A Flexible, Model-Based Approach to Partitioning Neural Variability. Neural Comput. 30, 1012– 1045 (2018).

20. Adibi, M., McDonald, J. S., Clifford, C. W. G. & Arabzadeh, E. Adaptation Improves Neural Coding Efficiency Despite Increasing Correlations in Variability. J. Neurosci. 33, 2108–2120 (2013).

21. Kheradpezhouh, E., Tang, M. F., Mattingley, J. B. & Arabzadeh, E. Enhanced Sensory Coding in Mouse Vibrissal and Visual Cortex through TRPA1. Cell Rep. 32, 107935 (2020).

22. Odagaki, Y., Kinoshita, M. & Toyoshima, R. Pharmacological characterization of M1 muscarinic acetylcholine receptor-mediated Gq activation in rat cerebral cortical and hippocampal membranes. Naunyn. Schmiedebergs. Arch. Pharmacol. 386, 937–947 (2013).

23. Dana, H. et al. High-performance calcium sensors for imaging activity in neuronal populations and microcompartments. Nat. Methods 16, 649–657 (2019).

24. Lee, C. C. Y., Kheradpezhouh, E., Diamond, M. E. & Arabzadeh, E. State-Dependent Changes in Perception and Coding in the Mouse Somatosensory Cortex. Cell Rep. 32, 108197 (2020).

25. Kohn, A., Coen-Cagli, R., Kanitscheider, I. & Pouget, A. Correlations and Neuronal Population Information. https://doi.org/10.1146/annurev-neuro-070815-013851 39, 237–256 (2016).

26. Polack, P. O., Friedman, J. & Golshani, P. Cellular mechanisms of brain state-dependent gain modulation in visual cortex. Nat. Neurosci. 16, 1331–1339 (2013).

27. Sabri, M. M. & Arabzadeh, E. Information processing across behavioral states: Modes of operation and population dynamics in rodent sensory cortex. Neuroscience 368, 214–228 (2018).

28. Marguet, S. L. & Harris, K. D. State-Dependent Representation of Amplitude-Modulated Noise Stimuli in Rat Auditory Cortex. J. Neurosci. 31, 6414–6420 (2011).

29. Pachitariu, M., Lyamzin, D. R., Sahani, M. & Lesica, N. A. State-dependent population coding in primary auditory cortex. J. Neurosci. 35, 2058–2073 (2015).

30. Reimer, J. et al. Pupil fluctuations track fast switching of cortical states during quiet wakefulness. Neuron 84, 355 (2014).

31. Eggermann, E., Kremer, Y., Crochet, S. & Petersen, C. C. H. Cholinergic Signals in Mouse Barrel Cortex during Active Whisker Sensing. Cell Rep. 9, 1654–1660 (2014).

32. Buzsaki, G. et al. Nucleus basalis and thalamic control of neocortical activity in the freely moving rat. J. Neurosci. 8, 4007–4026 (1988).

33. Metherate, R., Cox, C. L. & Ashe, J. H. Cellular bases of neocortical activation: Modulation of neural oscillations by the nucleus basalis and endogenous acetylcholine. J. Neurosci. 12, 4701–4711 (1992).

34. Do, J. P. et al. Cell type-specific long-range connections of basal forebrain circuit. Elife 5, (2016).

35. Staiger, J. F. & Petersen, C. C. H. Neuronal Circuits in Barrel Cortex for Whisker Sensory Perception. Physiol. Rev. 101, 353–415 (2021).

36. Feldmeyer, D., Qi, G., Emmenegger, V. & Staiger, J. F. Inhibitory interneurons and their circuit motifs in the many layers of the barrel cortex. Neuroscience 368, 132–151 (2018).

37. Lee, C. C. Y., Diamond, M. E. & Arabzadeh, E. Sensory Prioritization in Rats: Behavioral Performance and Neuronal Correlates. J. Neurosci. 36, 3243 LP – 3253 (2016).

38. Ohshiro, T., Angelaki, D. E. & Deangelis, G. C. A normalization model of multisensory integration. Nat. Neurosci. 14, 775–782 (2011).

39. Salinas, E. & Thier, P. Gain modulation: a major computational principle of the central nervous system. Neuron 27, 15–21 (2000).

40. Soma, S., Shimegi, S., Suematsu, N. & Sato, H. Cholinergic modulation of response gain in the rat primary visual cortex. Sci. Rep. 3, 1–7 (2013).

41. Ferguson, K. A. & Cardin, J. A. Mechanisms underlying gain modulation in the cortex. Nat. Rev. Neurosci. 2020 212 21, 80–92 (2020).

42. Thiele, A. & Bellgrove, M. A. Neuromodulation of Attention. Neuron 97, 769–785 (2018).

43. Pinto, L. et al. Fast modulation of visual perception by basal forebrain cholinergic neurons. Nat. Neurosci. 16, 1857–1863 (2013).

44. Beaman, C. B., Eagleman, S. L. & Dragoi, V. Sensory coding accuracy and perceptual performance are improved during the desynchronized cortical state. Nat. Commun. 8, (2017).

45. Engel, T. A. et al. Selective modulation of cortical state during spatial attention. Science 354, 1140–1144 (2016).

46. Khateb, M., Schiller, J. & Schiller, Y. State-Dependent Synchrony and Functional Connectivity in the Primary and Secondary Whisker Somatosensory Cortices. Front. Syst. Neurosci. 0, 94 (2021).

47. Middleton, J. W., Omar, C., Doiron, B. & Simons, D. J. Neural Correlation Is Stimulus Modulated by Feedforward Inhibitory Circuitry. J. Neurosci. 32, 506 (2012).

48. Jang, H. J. et al. Distinct roles of parvalbumin and somatostatin interneurons in gating the synchronization of spike times in the neocortex. Sci. Adv 6, 5333–5355 (2020).

49. Chen, N., Sugihara, H. & Sur, M. An acetylcholine-activated microcircuit drives temporal dynamics of cortical activity. Nat. Neurosci. 18, 892–902 (2015).

50. Cea-Del Rio, C. A., Lawrence, J. J., Erdelyi, F., Szabo, G. & Mcbain, C. J. Cholinergic modulation amplifies the intrinsic oscillatory properties of CA1 hippocampal cholecystokinin-positive interneurons. J. Physiol. 589, 609–627 (2011).

51. Pafundo, D. E., Miyamae, T., Lewis, D. A. & Gonzalez-Burgos, G. Cholinergic modulation of neuronal excitability and recurrent excitation-inhibition in prefrontal cortex circuits: Implications for gamma oscillations. J. Physiol. 591, 4725–4748 (2013).

52. Fazlali, Z., Ranjbar-Slamloo, Y., Adibi, M. & Arabzadeh, E. Correlation between cortical state and locus coeruleus activity: Implications for sensory coding in rat barrel cortex. Front. Neural Circuits 10, 14 (2016).

53. Matteucci, G. et al. Cortical sensory processing across motivational states during goal-directed behavior. Neuron 0, (2022).

54. Shirey, J. K. et al. A selective allosteric potentiator of the M1 muscarinic acetylcholine receptor increases activity of medial prefrontal cortical neurons and restores impairments in reversal learning. J. Neurosci. 29, 14271–14286 (2009).

55. James, N., Gritton, H., Kopell, N., Sen, K. & Han, X. Muscarinic receptors regulate auditory and prefrontal cortical communication during auditory processing. Muscarinic Recept. Regul. Audit. prefrontal cortical Commun. Dur. Audit. Process. 285601 (2018) doi:10.1101/285601.

56. Niell, C. M. & Stryker, M. P. Modulation of visual responses by behavioral state in mouse visual cortex. Neuron 65, 472 (2010).

57. Poulet, J. F. A. & Petersen, C. C. H. Internal brain state regulates membrane potential synchrony in barrel cortex of behaving mice. Nature 454, 881–885 (2008).

58. Diamond, M. E., Von Heimendahl, M., Knutsen, P. M., Kleinfeld, D. & Ahissar, E. ‘Where’ and ‘what’ in the whisker sensorimotor system. Nat. Rev. Neurosci. 9, 601– 612 (2008).

59. Zuo, Y., Perkon, I. & Diamond, M. E. Whisking and whisker kinematics during a texture classification task. Philos. Trans. R. Soc. Lond. B. Biol. Sci. 366, 3058–3069 (2011).

60. Adibi, M., Diamond, M. E. & Arabzadeh, E. Behavioral study of whisker-mediated vibration sensation in rats. Proc. Natl. Acad. Sci. U. S. A. 109, 971–976 (2012).

61. Fassihi, A., Akrami, A., Esmaeili, V. & Diamond, M. E. Tactile perception and working memory in rats and humans. Proc. Natl. Acad. Sci. U. S. A. 111, 2331–2336 (2014).

62. Yang, H., Kwon, S. E., Severson, K. S. & O’Connor, D. H. Origins of choice-related activity in mouse somatosensory cortex. Nat. Neurosci. 19, 127–134 (2015).

63. O’Connor, D. H., Peron, S. P., Huber, D. & Svoboda, K. Neural activity in barrel cortex underlying vibrissa-based object localization in mice. Neuron 67, 1048–1061 (2010).

64. Gould, R. W. et al. Role for the M1 Muscarinic Acetylcholine Receptor in Top-Down Cognitive Processing Using a Touchscreen Visual Discrimination Task in Mice. ACS Chem. Neurosci. 6, 1683–1695 (2015).

65. Yousuf, H., Girardi, E. M., Crouse, R. B. & Picciotto, M. R. Muscarinic antagonists impair multiple aspects of operant discrimination learning and performance. Neurosci. Lett. 794, 137025 (2023).

66. Gasselin, C., Hohl, B., Vernet, A., Crochet, S. & Petersen, C. C. H. Cell-type-specific nicotinic input disinhibits mouse barrel cortex during active sensing. Neuron 109, 778–787.e3 (2021).

67. Chen, N., Sugihara, H. & Sur, M. An acetylcholine-activated microcircuit drives temporal dynamics of cortical activity. Nat. Neurosci. 2015 186 18, 892–902 (2015).

68. Gainey, M. A., Aman, J. W. & Feldman, D. E. Rapid Disinhibition by Adjustment of PV Intrinsic Excitability during Whisker Map Plasticity in Mouse S1. J. Neurosci. 38, 4749–4761 (2018).

69. Lee, S., Kruglikov, I., Huang, Z. J., Fishell, G. & Rudy, B. A disinhibitory circuit mediates motor integration in the somatosensory cortex. Nat. Neurosci. 2013 1611 16, 1662–1670 (2013).

70. Zagha, E. & McCormick, D. A. Neural Control of Brain State. Curr. Opin. Neurobiol. 0, 178 (2014).

71. Joshi, S. & Hawken, M. J. Loose-patch-juxtacellular recording in vivo-A method for functional characterization and labeling of neurons in macaque V1. J. Neurosci. Methods 156, 37–49 (2006).

72. Pinault, D. The juxtacellular recording-labeling technique. Neuromethods vol. 54 41– 75 (2011).

73. Carandini, M. & Churchland, A. K. Probing perceptual decisions in rodents. Nat. Neurosci. 16, 824 (2013).

